# Chromatographic, Immuno-inflammatory, and nephrotoxic Assessment of selected culinary oils in Experimental Models

**DOI:** 10.1101/2023.08.30.555600

**Authors:** Jude Ogechukwu Okoye, Chiagoziem Moral Delu-Mozie, Maureen Ugochukwu Nwachioma, Uchenna Benjamin Modozie

## Abstract

**Background:** Dietary oils are crucial for everyday human nutrition. They contain essential fatty acids and support a range of physiological functions. However, concerns regarding their possible toxicity have been expressed, particularly concerning the elevation of cholesterol levels, particularly Low-Density Lipoprotein (LDL). This study investigated the composition of selected dietary oils and determined their physiologic effects and the micro-architectural integrity of the kidney.

**Methods:** In this experimental study, 30 Albino rats were used in this study. The animals were divided into 5 groups: Groups A, B, C, D, and E (n=6 each). Group A (control) received normal rat pellets only while Groups B, C, D, and E received rat pellets mixed with Avocado oil, Coconut oil, Palm oil, and Peanut oil. Blood samples were drawn, and kidneys were excised. Data generated from biochemical, haematological, and histological investigations were analyzed using ANOVA, Pearson’s correlation, and *Post hoc* test. Significance was set at p< 0.05.

**Results:** The results indicated significant differences in fatty acid levels between oils (p< 0.05). Higher levels of oleic acid, lauric acid, palmitic acid, and oleic acid were found in coconut oil, Palm oil, Avocado oil, and Peanut oil, respectively. Significant differences in urea levels were observed between the control group and other treatment groups (p= 0.001). Group B had lower levels of triglyceride while groups C and D had higher levels of LDL and organ weight, respectively compared with the control group (p= 0.035, 0.042, and 0.008, respectively). Group E had a higher neutrophil-lymphocyte ratio, mean corpuscular haemoglobin concentration, lower lymphocyte-monocyte ratio, and red cell distribution width (p= 0.325, 0.025, 0.068, and 0.053, respectively). Kidney sections revealed varying degrees of necrosis and inflammation,

**Conclusion:** The study provides valuable insights indicating potential oil-induced effects on health. It advises caution during the application of the oils in culinary activities.

## Introduction

Dietary oils, including palm, avocado, peanut, and coconut oils, are crucial for human nutrition. They contain essential fatty acids and support a range of physiological functions, highlighting their importance due to their distinctive compositions and potential health advantages. However, concerns regarding their possible toxicity have been expressed, particularly concerning the elevation of Low-Density Lipoprotein (LDL) cholesterol levels [1]. Unlike the larger lipoprotein particles such as LDL and Triglyceride, increasing concentrations of High-Density Lipoprotein (HDL) particles are associated with decreasing accumulation of atherosclerosis and macrophages within the walls of arteries, reducing the risk of sudden plaque ruptures, cardiovascular disease, stroke, and other vascular diseases [2]. Palm oil, extracted from the fruit of *Elaeis guineensis jacquin*) has sparked debates about its effects on cardiovascular health due to its saturated and unsaturated fatty acids composition. It also contains tocotrienols and carotenoids that may have antioxidant properties [3,4]. It is also rich in vitamin E, which inhibits cholesterol synthesis and protects the cell membrane from lipid peroxidation, thus conferring a protective effect against cardiovascular and neurodegenerative diseases and supporting the treatment of cancer [5]. The high vitamin E content in Pal has been shown to have anti-inflammatory effects in myocardial tissue in LPS-induced sepsis models [6]. Avocado oil, extracted from the flesh of *Persea americana*, is rich in monounsaturated fatty acids, especially oleic acid. It has potential anti-inflammatory and can improve lipid profiles. However, overconsumption can lead to weight gain due to its high-calorie content [7]. Peanut oil, derived from *Arachis hypogaea*, has a balanced ratio of mono- and polyunsaturated fatty acids, and a moderate amount of saturated fat. It has been suggested that it may have potential benefits for cardiovascular health [8]. Coconut oil, derived from the meat of the *Cocos nucifera*, is high in saturated fat. It has potential health benefits, including antimicrobial effects and weight management due to its medium-chain fatty acids [1]. Free fatty acids (FFAs) are carboxylic acids with long saturated or unsaturated aliphatic chains. They are important energy sources in most body tissues and show critical functions such as receptor signaling, gene expression, and systemic fuel energy homeostasis regulation under various physiological conditions [9-11]. Essential FFAs, such as linoleic acid (LA; omega-6 FAs) and alpha-linolenic acid (ALA; ome-ga-3 FAs), are plant-based and necessary for health [12-14]. ALA and LA can be converted to longer-chain omega-3 FFAs such as eicosapentaenoic acid (EPA) and docosahexaenoic acid (DHA) as well as longer-chain omega-6 FAs such as arachidonic acid [15]. This study offers insights into making informed dietary choices by assessing the nutritional compositions, bioactive compounds, and health effects of palm oil, avocado oil, Peanut oil, and coconut oil in a balanced evaluation of available evidence.

## Materials and Methods

### Oil extraction and Free Fatty Acid Detection

Coconut, Peanut, Avocado, and Palm oils were extracted by the Dry heat extraction method, Heat dependent physical-mechanical pressing method, Low heat Air dry pressing method, and Hot-pressing method, respectively [4]. The FFA in Palm fruits, Coconut meat, Peanut seed, and Avocado pear were extracted using the rapid and highly sensitive gas chromatographic method and flame ionization detector (Column: Restek 15 Meter MXT-1; Carrier: Helium AT 5 PSI) as described by Azeman et al. and de Vasconcelos et al. [16,17]. The FFAs are separated by the interaction between the stationary and mobile phases along the column before being detected by the detector.

### Animal and plant handling

The plants (Avocado pear, Coconut seed, Palm fruit, and Peanut were collected from Nnewi; latitude 5.976 and longitude 6.895. The plant oil was extracted accordingly, and the study was carried out at the Animal House Research Area, domiciled in the Physiology Department of Nnamdi Azikiwe College of Health Science and Technology, Okofia, Nnewi, Anambra State, Nigeria. The animals were fed with rat chow pellets and libitum with a normo-lipidemic and hyper-lipidemic diet, as well as water was made available to the rats in water bottles of the downspout type (drinking nozzle facing downward). The animals were fed with standard Pfizer-branded rodent feed from Livestock Feed, Nigeria Ltd. The feed is composed of 21% protein, 2% fat, 12.8% Can starch, 4.2% Salt and vitamins mixture and vitamins, and water. They were maintained under a (12h light and 12h dark photo cycle) throughout the experiment. The animals were weighed before, during, and after the research to ensure standard as well as monitor the pattern of the changes in weight during the experiment.

### Research design

A total of 30 Albino rats, weighing (120 to 140 grams), were used in this study. The animals were divided into 5 groups, Groups A, B, C, D, and E (n=6 each). Group A (control) was fed with rat pellets and water only. Group B was fed with rat pellets (10 grams) mixed with avocado oil (English pear oil), Group C was fed with rat pellets (10 grams) mixed with coconut oil, and Group D was fed with rat pellets (10 grams) mixed with Palm oil, while Group E was fed with Rat pellets (10 grams) mixed with Peanut-oil. The lethal dose (LD50) and effective dose of the extracted oils were approximately 5000mg/kg and 2500 mg/kg, respectively (Ahmad et al, 2016). The research lasted for 28 days. At the end of the experiment, after 12 hours of fasting, the albino rats were sacrificed, and blood samples were collected via retro-orbital sinus plexus from each albino rat in the control and treatment groups in EDTA anticoagulant sample bottles. Blood samples were left to clot and centrifuged at 5000 rpm for 10 minutes to separate the serum. Following blood collection, the biological models were decapitated for blood collection. Following laparotomy, the kidneys were excised (with the aid of a sharp scalpel blade), weighed, and fixed in 10% formalin immediately.

### Sample handling

Full blood count was assessed using an Automated haematology analyzer (Sysmex XN-450/XN-430; CLSI Procedure. This device analyzed blood using hydrodynamic impedance, flow cytometry with a semiconductor laser, SLS-hemoglobin, and RBC pulse height detection. The lipid profile includes total cholesterol (TC), high-density lipoproteins (HDL), Low-density lipoproteins (LDL), and Triglycerides (TG). The animals’ fasting lipid profile was carried out using the enzymatic method as described by Ochei et al. [18]. The total white cell count (TWBC) (neutrophil-to-lymphocyte ratio (NLR), platelet-to-lymphocyte ratio (PLR), Lymphocyte-to-monocyte ratio (, and Platelets-neutrophils to lymphocytes ratio (PNLR) were calculated for the subgroups.

### Statistical analysis

Data was analyzed using the Statistical Package for Social Sciences, version 21 computer software (SPSS). Analysis of Variance (ANOVA) and Student’
ss t-test was used to analyze the data. The experimental data were expressed as the mean ± standard error of mean. The limit for statistical significance was set at p ≤ 0.05.

## Results

### Free fatty acid analysis

The loss of two animals in groups C, D, and E suggests that some of the fatty acids in coconut, Palm, and Peanut oils have some detrimental effects on animal health (Figure 1). Therefore, blood parameters were examined.

**Figure 1:**
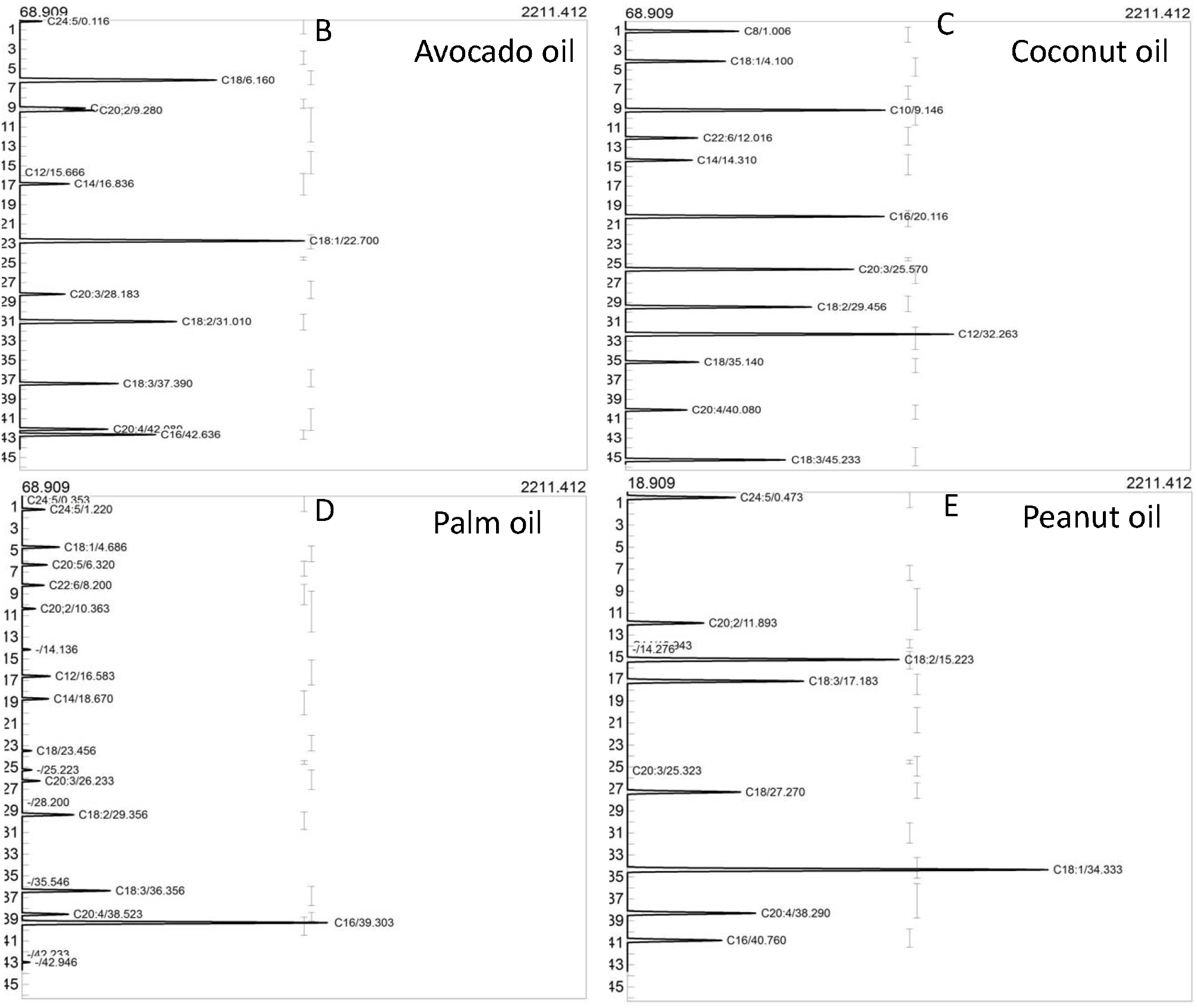
Gas chromatographic analysis of FFA in some culinary oils. In Figure 1, the percentage of lauric acid (saturated fatty acid) was higher in Coconut oil compared with Palm Oil, Avocado oil, and Peanut oil (p< 0.001). The percentages of Myristic and Palmitic acids (saturated fatty acids) were higher in Palm oil compared with Coconut Oil, Avocado oil, and Peanut oil at p< 0.001 (Figure 1). Higher percentages of stearic acid (saturated fatty acid), Linolenic acid (monounsaturated fatty acid), and Docosapentaenoic and Docosahexaenoic acids (polyunsaturated fatty acid) were found in Avocado oil than in Palm oil, coconut oil, and Peanut oil (p< 0.001). The percentages of Oleic and Linoleic acids (monounsaturated fatty acids), Eicosapentaenoic, Arachidonic, and Lignoceric acids (Polyunsaturated fatty acids) were higher in Peanut oil compared with Avocado oil, Coconut Oil, and Palm oil (p< 0.001).

In terms of Urea conc., there were significant differences between group A and groups C, D, and E (at p= 0.001, 0.005, 0.021, respectively). There were also significant differences between group B and groups C, D, and E (at p= 0.001, 0.004, 0.001, respectively). In terms of R.O.W.K, there were significant differences between groups A and D, and groups B and D, groups C and D at p= 0.008, 0.045, 0.019 (Table 2). Overall, the findings suggest that the administration of different oils (Avocado, Coconut, Palm, and Peanut) did not significantly alter the overall electrolyte levels, Significant direct relationships were observed between Potassium and Sodium, The data in Figure 2 reveals varying patterns of significance across different parameters and experimental groups. While some parameters, especially mean corpuscular haemoglobin concentration (MCHC) and red cell distribution width-standard deviation (RDW-SD), show significant differences among the groups, others demonstrate no significant variations. The groups treated with Peanut oil had higher and lower MCHC and RDW-SD, respectively compared with other treatment groups (*p* ≤ 0.05). These findings suggest that the experimental treatments may have influenced specific blood characteristics. Regarding immune-inflammatory assessment, group B had a higher PLR and PNLR following treatment compared with other groups (p> 0.05) while group E had a higher NLR but lower PLR and LMR compared with other groups (p> 0.05).

**Table 1a:** Results for fatty acid composition in Coconut and Avocado Oil.

| S/N | Components | Compounds | Coconut Oil Conc. |  | Avocado Oil Conc. |  |
| --- | --- | --- | --- | --- | --- | --- |
|  |  |  | ppm | % | ppm | % |
| 1 | C8 | Caprylic acid | 6.9049 | 7.399 | UD | UD |
| 2 | C10 | Capric acids | 8.1433 | 8.726 | UD | UD |
| 3 | C12 | Lauric acid | 33.7234 | 36.135 | 0.0199 | 0.025 |
| 4 | C14 | Myristic acid | 6.5784 | 7.049 | 5.1557 | 6.364 |
| 5 | C16 | Palmitic acid | 8.3251 | 8.920 | 11.0548 | 13.645 |
| 6 | C18 | Stearic acid | 7.6552 | 8.203 | 11.5194 | 14.219 |
| 7 | C18:1 | Oleic acid | 4.0822 | 4.374 | 34.8725 | 43.045 |
| 8 | C18:2 | Linoleic acid | 3.2739 | 3.508 | 3.2781 | 4.046 |
| 9 | C18:3 | Linolenic acid | 4.1621 | 4.461 | 2.6763 | 3.304 |
| 10 | C20:2 | Docosapentaenoic acid | UD | UD | 2.6744 | 3.301 |
| 11 | C20:3 | Eicosapentaenoic acid | 2.1807 | 2.337 | 0.5464 | 0.675 |
| 12 | C20:4 | Arachidonic acid | 2.6190 | 2.806 | 3.4235 | 4.226 |
| 13 | C22:6 | Docosehaxaenoic acid | UD | UD | 4.9914 | 6.161 |
| 14 | C24:5 | Lignoceric acid | UD | UD | 0.8020 | 0.991 |
| Total |  |  | 93.3275 | 100 | 81.0146 | 100 |

Keys: UD= Undetected

**Table 1b:** Fatty acid composition in Palm and Peanut Oil.

| S/N | Components | Compound | Palm Oil Conc. |  | Peanut Oil Conc. |  |
| --- | --- | --- | --- | --- | --- | --- |
|  |  |  | ppm | % | ppm | % |
| 1 | C12 | Lauric acid | 4.3327 | 9.247 | UD | UD |
| 2 | C14 | Myristic acid | 3.4941 | 7.446 | 0.0900 | 0.001 |
| 3 | C16 | Palmitic acid | 24.1593 | 51.564 | 3.3207 | 3.951 |
| 4 | C18 | Stearic acid | 2.3342 | 4.982 | 4.0475 | 4.814 |
| 5 | C18:1 | Oleic acid | 1.8959 | 4.047 | 46.4656 | 55.265 |
| 6 | C18:2 | Linoleic acid | 1.1828 | 2.525 | 12.1012 | 14.393 |
| 7 | C18:3 | Linolenic acid | 2.4639 | 5.259 | 4.3067 | 5.122 |
| 8 | C20:2 | Docosapentaenoic acid | 0.9005 | 1.922 | 1.8585 | 2.210 |
| 9 | C20:3 | Eicosapentaenoic acid | 0.3131 | 0.668 | 0.0120 | 0.014 |
| 10 | C20:4 | Arachidonic acid | 2.1130 | 4.510 | 8.5186 | 10.132 |
| 11 | C22:6 | Docosehaxaenoic acid | 2.4835 | 5.301 | UD | UD |
| 12 | C24:5 | Lignoceric acid | 1.1798 | 2.518 | 3.3575 | 3.993 |
| Total |  |  | 46.8528 | 100 | 84.0783 | 100 |

Keys: UD= Undetected

**Table 2:** Mean comparison of electrolyte levels across the experiment groups.

| Groups | N<sup>o</sup> of | Mean ± Standard Deviation (SD) | | | | | |
| --- | --- | --- | --- | --- | --- | --- | --- |
|  | Animal | Potassium | Sodium | Chloride | HCO | Creatinine | Urea |
| | | mmol/L | mmol/L | mmol/L | mmol/L | μmol/L | μmol/L |
| **Group A** | 6 | 4.93±0.46 | 142.00±2.61 | 101.50±1.38 | 24.17±1.17 | 51.83±2.96 | 12.68±0.71 |
| **Group B** | 5 | 4.72±0.50 | 142.40±2.70 | 102.60±1.67 | 23.80±1.30 | 66.20±3.01 | 13.52±2.70 |
| **Group C** | 4 | 4.75±0.62 | 141.75±2.75 | 101.50±2.38 | 25.00±1.41 | 61.25±1.78 | 9.61 ±0.81 |
| **Group D** | 4 | 4.80±0.37 | 143.75±2.63 | 103.25±2.36 | 24.25±0.96 | 68.25±2.45 | 10.28±0.79 |
| **Group E** | 4 | 5.08±0.35 | 143.25±1.26 | 102.50±2.08 | 24.75±0.96 | 45.25±1.12 | 9.61 ±1.09 |
| F-value |  | 0.427 | 0.493 | 0.698 | 0.731 | 1.140 | 7.244 |
| P-value |  | 0.787 | 0.741 | 0.603 | 0.582 | 0.369 | 0.001* |
| **A vs B** |  | 0.463 | 0.794 | 0.360 | 0.615 | 0.232 | 0.358 |
| **A vs C** |  | 0.553 | 0.878 | 0.999 | 0.289 | 0.456 | 0.005* |
| **A vs D** |  | 0.666 | 0.290 | 0.178 | 0.914 | 0.201 | 0.021* |
| **A vs E** |  | 0.646 | 0.447 | 0.434 | 0.454 | 0.601 | 0.005* |

\*Significance is set at $p \leq 0.05$ . Statistical analysis: ANOVA, *Post hoc* Test. Keys: Bicarbonate= HCO, Group A= Control, Group B= Avocado oil, Group C= Coconut oil, Group D= Palm Oil, Group E= Peanut Oil

**Table 3:** Comparison of lipid profile and relative organ weight.

| Groups | N<sup>o</sup> of | Mean ± Standard Deviation (SD) | | | | |
| --- | --- | --- | --- | --- | --- | --- |
|  | Animal | TC | TG | HDL | LDL | R.O.W.K |
|  |  | mg/dL | mg/dL | mmol/L | mmol/L |  |
| **Group A** | 6 | 2.02±0.22 | 0.87±0.34 | 0.64±0.15 | 0.98±0.10 | 0.69±0.18 |
| **Group B** | 5 | 2.34±0.50 | 0.60±0.07 | 0.73±0.18 | 1.33±0.32 | 0.75±0.04 |
| **Group C** | 4 | 2.58±0.49 | 0.66±0.07 | 0.69±0.55 | 1.60±0.47 | 0.71±0.05 |
| **Group D** | 4 | 2.05±0.70 | 0.69±0.14 | 0.53±0.07 | 1.21±0.63 | 0.92±0.10 |
| **Group E** | 4 | 2.03±0.65 | 0.55±0.06 | 0.69±0.13 | 1.08±0.59 | 0.82±0.10 |
| F-value |  | 1.038 | 2.107 | 0.410 | 1.390 | 2.815 |
| P-value |  | 0.415 | 0.122 | 0.799 | 0.277 | 0.056 |
| **A vs B** |  | 0.314 | 0.035* | 0.550 | 0.203 | 0.423 |
| **A vs C** |  | 0.104 | 0.106 | 0.772 | 0.042* | 0.850 |
| **A vs D** |  | 0.917 | 0.173 | 0.507 | 0.429 | 0.008* |
| **A vs E** |  | 0.976 | 0.020* | 0.750 | 0.734 | 0.103 |

\*Significance is set at $p \leq 0.05$ . Statistical analysis: ANOVA, *Post hoc* Test. Keys: R.O.W.K= Relative organ weight of Kidney, Group A= Control, Group B= Avocado oil, Group C= Coconut oil, Group D= Palm Oil, Group E= Peanut Oil

**Figure 2:**
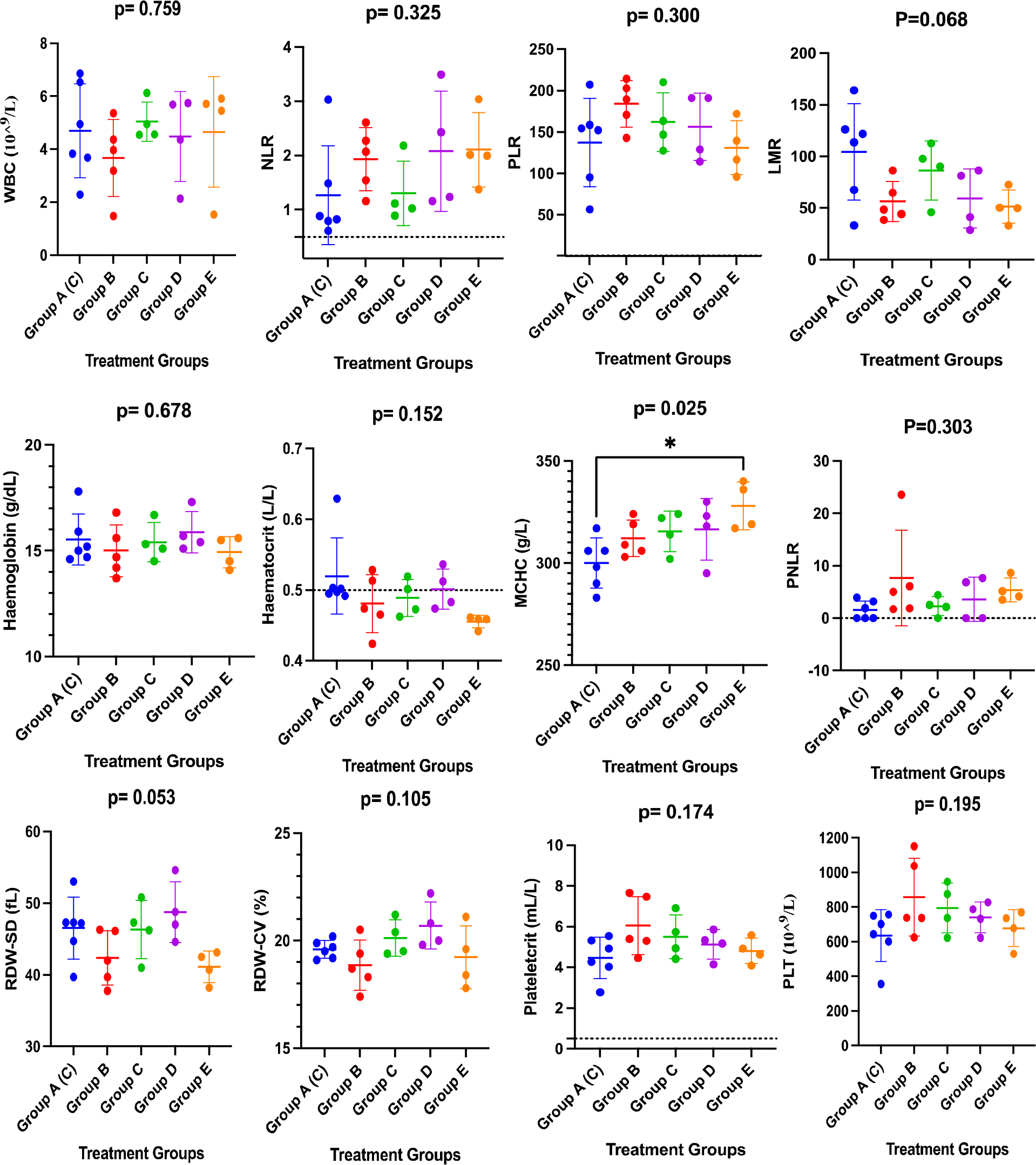
Graphical presentation of blood parameters across experimental groups.

**Figure 2:**
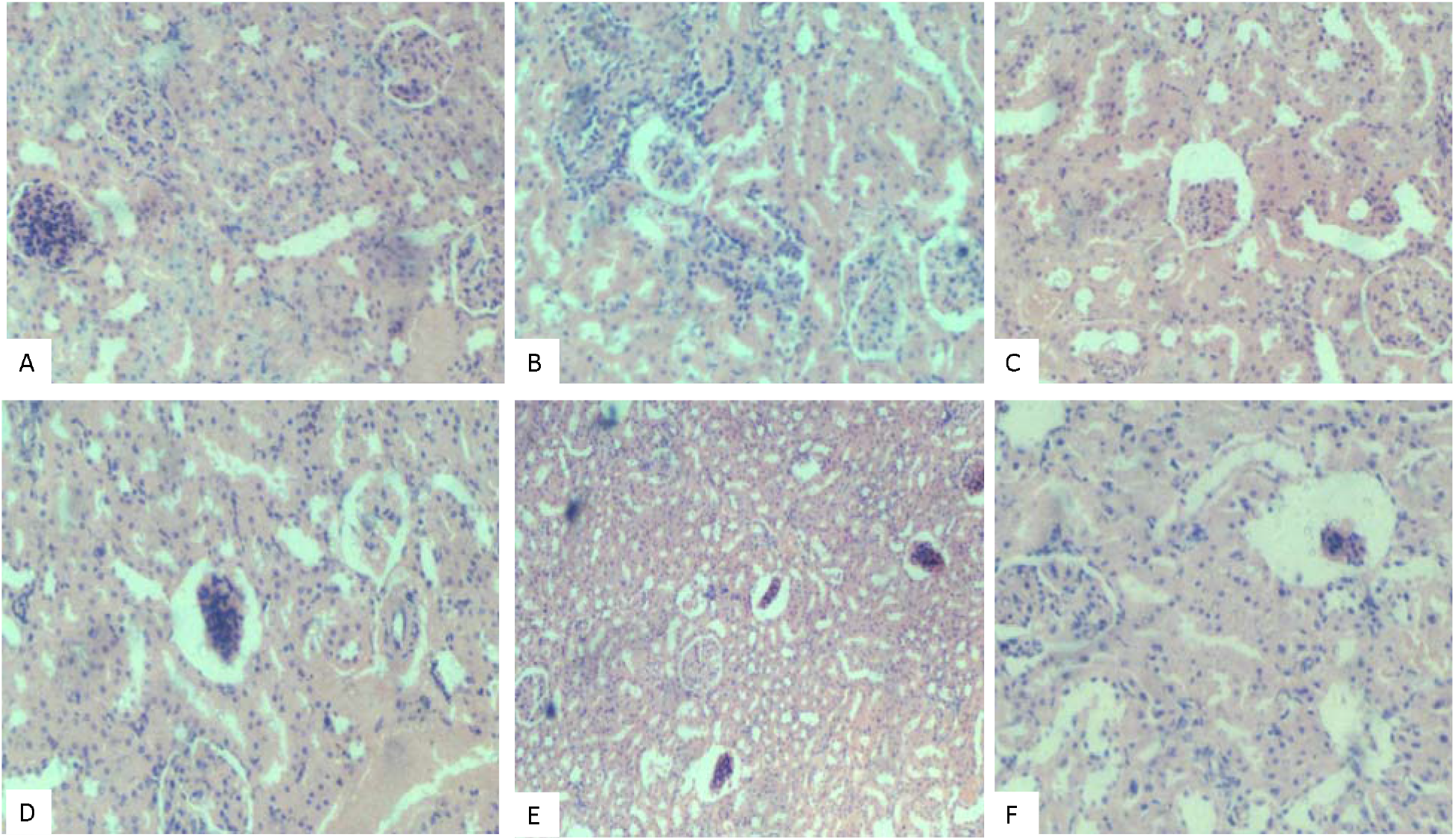
Photomicrograph of kidney sections stained by H&E technique.

Potassium and Chloride, Sodium and Chloride, Sodium and Bicarbonate, Creatinine, and R.O.W.K at p= 0.004, 0.021, 0.000, 0.014, and 0.000, respectively while significant inverse relationships were observed between Urea and Bicarbonate at p= 0.010. There were significant direct relationships between BRC and TG, HGB and TG, HCT and TG, RDW-CV, and LDL at p= 0.049, 0.021, 0.004 and 0.018, respectively. These findings highlight potential variations in lipid profiles and relative organ weight among the different oil-treated groups. The variations in TG and LDL levels suggest that the types of oils administered could impact lipid metabolism. In the pairwise comparisons (Figure 2), there were significant differences observed in certain parameters such as platelet count (A vs. B; p= 0.029), RDW-CV (B vs. D, and D vs. E; p= 0.014 and 0.054, respectively), HCT (A vs E; p= 0.017), MCV (D vs E; p= 0.044), MCHC (A vs D and A vs E; p= 0.043 and 0.002, respectively), and RDW-SD (A vs E, B vs D, D vs E; p= 0.044, 0.025, and 0.012, respectively). A higher platelet mass was observed in group B (6.05 ± 1.42) compared with the control group (4.47 ± 1.01) at p= 0.022.

Figure 2A shows apparently healthy glomeruli. Figure 2B shows evidence of mild necrosis and inflammation with marked interstitial infiltration of inflammatory cells. Figure 2C shows atrophic glomeruli and moderate necrosis. Figure 2D: shows atrophic glomeruli and moderate necrosis. Figure 2E shows extensive glomerulo-atrophy (X100). Figure 2F shows atrophic glomeruli and mild necrosis (X400). The architectural changes in Figure 2E are consistent with glomerulonephritis.

## Discussion

The present study investigated the FFA composition of Coconut oil, Palm oil, Avocado oil, and Peanut oil, and assessed their potential effects on physiological parameters. According to the FFA analysis, variations in fatty acid composition were identified among different oils. For instance, coconut oil contained higher levels of lauric acid, while avocado oil demonstrated a greater concentration of stearic acid, linolenic acid, and other polyunsaturated fatty acids. This suggests that avocado oil has the potential to serve as a valuable source of diverse fatty acids with health benefits. Similarly, Peanut oil was found to have a unique fatty acid profile, with higher proportions of oleic and linoleic acids, eicosapentaenoic, arachidonic, and lignoceric acids. The FFA composition may have implications for their nutritional and health effects. While there were no significant alterations in overall electrolyte levels, some specific relationships were noted. Notably, significant direct relationships were observed between potassium and sodium, potassium and chloride, sodium and chloride, sodium and bicarbonate, creatinine, and the relative weight of the kidney. These findings suggest intricate interplays between these parameters that may be influenced by the oils’ compositions. The variation in lipid profile analysis suggests that the oils may have varying effects on lipid metabolism, which could contribute to their potential cardiovascular implications. These omega-3 and omega-6 FFAs are precursors of eicosanoid inflammatory mediators such as leukotrienes (LTs), prostaglandins (PGs), thromboxanes (TXs), and resolvins [14].

There was an abundance of oleic acid, an omega-9 fatty acid, in the peanut oil used in this study. Apart from the potential of reducing the risk of hypertension, Oleic acid-containing diets are believed to modulate immune response towards eliminating pathogenic microbes [19]. However, a higher arachidonic acid, an omega-6 FFA, was observed in peanut oil compared with other oils. Studies have shown that a high level of arachidonic acid contributes to thrombosis, inflammation, atherosclerosis, obesity, and diabetes [12,14]. Evidence such as decreased RDW-SD and high MCHC suggest anemia, which aligns with the histological findings. Although there was a lower PLR among animal models that received a peanut-enriched diet (group E), a lower LMR was also compared with other groups including the control. A low LMR reflects an active inflammation status and has been associated with worse overall survival and disease-free survival [20]. Studies have shown that NLR, PLR, and PNLR could be used as prognostic biomarkers based on their potential to predict poor overall survival and unfavourable progression-free survival [21,22]. The high NLR and PNLR may have contributed to the death of some animals fed the peanut-enriched diet. It could be argued that the presence of arachidonic acid in peanut oil counteracts the oleic acid content under physiological conditions.

This study revealed high polyunsaturated fatty acids (omega-3 FFA; Linolenic acid, Docosapentaenoic acid, and Eicosapentaenoic acid) in Avocado oil. Studies show that omega-3 FFA reduces the incidence of coronary heart disease [12-15]. The high PLR and PNLR among animals fed an Avocado-enriched diet suggests that other FFA components of the oil may be exerting counteracting effects on the omega-3 FFA. The high plateletcrit and platelet count in group B suggests that the FFA in Avocado oil may have induced thrombocytosis. This increases the risk of thrombosis and death. Coconut oil’s high lauric acid content may offer antimicrobial properties [1], but its saturated fat content has raised concerns regarding its impact on cardiovascular health. This study shows a high amount of cholesterol and LDL and a lower amount of sodium and chloride among models fed with coconut oil compared with other treatment groups. The latter might be an explanation for the atrophic glomeruli and nephro-necrosis. An earlier study found that consumption of coconut oil can lead to negative health effects such as metabolic dysfunctions, adipose inflammation, and hepatic lipid accumulation [2]. Thus, caution is advised when consuming coconut oil, as excessive intake can result in renal failure despite their potential benefits for reducing fatigue and improving growth [14].

In this study, palm oil contained higher levels of omega-3 fatty acids than other oils. However, the animal models fed a palm oil-enriched diet had lower HDL and creatinine levels and higher organ weight than the control group. This correlates well with the histological findings. The observed toxicity could be attributed to the high content of palmitic acid, a saturated fatty acid, in Palm oil. Literature shows that palmitic acid induces dyslipidemia, hyperglycemia, fat accumulation, and pro-inflammatory effects via toll-like receptor 4 (Carta et al., 2017). It is also believed to regulate disease development and progression of metabolic diseases and cancer (Fatima et al., 2019). However, it’s important to note that this study has limitations. Further research is needed to investigate the individual effects of these oils on biomarkers of health, lipid metabolism, and chronic disease risk. Long-term studies that consider the oils’ effects on actual health outcomes will provide more conclusive insights into their nutritional significance.

## Conclusion

This study provides valuable insights into the free fatty acid compositions of Coconut oil, Palm oil, Avocado oil, and Peanut oil, shedding light on their potential health effects. The variations in FFA profiles suggest distinct nutritional profiles for each oil, which could influence their physiological and health implications. The observed effects on physiological parameters such as urea concentration and kidney weight highlight the importance of considering the oils’ potential renal impact. Similarly, the lipid profile variations underscore the oils’ potential cardiovascular effects.

## Ethical Declarations

The ethical approval for this research was obtained from the College of Health Sciences Ethics Committee (NAU/FHST/2021/MLS47). The principles of laboratory animal care were followed as described by the National Institute of Health (NIH publication number 85-23, revised 1985). All experiments involving the adult Albino rats were performed according to the guide of the National Research Council (2011).

## Acknowledgment

Authors owe a great deal of gratitude to the staff of the Springboard Research Laboratory Awka for their technical assistance.

## Author Contributions Statement

J.O.O. conceptualized and designed the study. Authors C.D. and M.O. carried out the experiment. UBM carried out the gas chromatographic analysis. J.O.O. carried out the statistics and wrote the first draft of the manuscript. All authors approved the final version of the manuscript.

## Data availability

The datasets generated during and/or analyzed during the current study are available from the corresponding author on reasonable request.

